# Birth order specified recruitment of motor circuits during spontaneous neural activity in zebrafish embryo

**DOI:** 10.1101/2024.08.27.609675

**Authors:** Maroun Abi Younes, Mohamad Rima, Jean-Pierre Coutanceau, Ninon Peysson, Fatemeh Hoseinkhani, Edson Rodrigues, Erika Bullier, Pascal Legendre, Jean-Marie Mangin, Elim Hong

## Abstract

Modular organization of spinal neural circuits control dynamic regulation of locomotion. However, it is unknown when or how the distinct microcircuits emerge during development. We carried out high-resolution calcium imaging of neural activity driving the first motor behavior in one day old zebrafish embryo. During this period, at least two waves of neurogenesis occur to generate primary and secondary motoneurons. We found that embryos first display a single highly synchronized rhythmic neuronal circuit containing interneurons and motoneurons. Later, two distinct interneuron-motoneuron circuits emerge with one containing early-born motoneurons displaying low-frequency activity and the other containing later-born motoneurons with high-frequency activity. The results indicate a mode of birth order determined microcircuits where neurons that are born together are recruited together. Nicotine affected neuronal activity frequency, revealing a functional role for cholinergic signaling in the emergence of patterned spinal circuits. Indeed, we found aberrant arrhythmic synchronized activity in mutants for *cholineacetyltransferase-a* where acetylcholine is no longer synthesized. Overall, we reveal the sequential recruitment of birth order specified microcircuits during the emergence of the earliest motor behavior and highlight a conserved role for cholinergic signaling in regulating rhythmic neural activity in the embryonic spinal cord.

## Introduction

Common to all animals is the ability to dynamically move throughout environment to seek shelter, forage for food, escape prey and find mates. A main focus of interest in the field of neuroscience is understanding how neural circuits control diverse motor behaviors. Studies in mammals and fish have identified distinct neural circuit modules underlying locomotion (El Manira, 2023; Hägglund et al., 2013; McLean et al., 2007). However, it is yet unclear when these circuits emerge during development.

Pharmacological, genetic, and optogenetic approaches combined with electrophysiological recordings and calcium imaging revealed the occurrence of synchronized spontaneous neural activity (SNA) across species in various developing circuits, including the spinal cord (Blankenship & Feller, 2010; Feller, 1999; Kirkby et al., 2013). SNA plays a role in the maturation of neural circuits as perturbing SNA during critical developmental periods in the spinal cord and other developing systems result in aberrant effects on axonal projection (Hanson & Landmesser, 2006; Kastanenka & Landmesser, 2010; Kirkby et al., 2013; Tezuka et al., 2022).

In the developing spinal cord of chick and mouse embryos, SNA is initiated when motoneurons start extending their axons toward the muscles (Hanson & Landmesser, 2003; O’Donovan & Landmesser, 1987). SNA consists of patterned recurring activity that progresses from synchronized activity in the whole spinal cord to alternating between the left and right sides (Momose-Sato & Sato, 2013; O’Donovan & Landmesser, 1987). The left-right rhythmic alternating activity is coordinated by a central pattern generator (CPG), which is thought to be the source of rhythmic and stereotyped behavior such as walking or swimming (Rancic & Gosgnach, 2021). SNA is initially dependent on cholinergic signaling in the spinal cord as blocking nicotinic acetylcholine receptors (nAChR) using pharmacology results in silencing SNA (Blankenship & Feller, 2010; Milner & Landmesser, 1999; Momose-Sato & Sato, 2020).

In the zebrafish spinal cord, SNA is first observed at 17 hours post fertilization (hpf), and is primarily dependent on gap junctions and calcium ion conductances (Saint-Amant & Drapeau, 2001; Warp et al., 2012). Calcium imaging shows that the activity begins in groups of neighboring cells, which become ipsilaterally correlated and contralaterally anti-correlated within a few hours (Wan et al., 2019; Warp et al., 2012).

The frequency of left-right alternating motor activity, referred to as coiling behavior, decreases after 19 hpf and stops around 30 hpf (Drapeau et al., 1999; Saint-Amant & Drapeau, 2000). Due to the rapid development of the zebrafish embryo, several rounds of neurogenesis occur within the first day (Alunni et al., 2013; Schmidt et al., 2013). However, how the different groups of sequentially born neurons incorporate into the spinal circuit during the earliest coiling motor behavior have not been investigated.

We performed high resolution calcium imaging of neurons in the rostral spinal cord of 1 day old zebrafish embryos co-expressing GCaMP in most post-mitotic spinal neurons and red fluorescent protein (RFP) in motoneurons [*Tg(elavl3:GCaMP6f;mnx1:Gal4;UAS:RFP)*] during SNA. We found that spinal neurons form a single functional circuit with synchronized rhythmic activity that alternate between the left and right sides of the spinal cord. In as little as a couple of hours later, two microcircuits emerge consisting of fast- or low-frequency activity in the spinal cord. Further analysis shows that primary motoneurons (CaP) with larger soma size mostly display low-frequency activity, whereas motoneurons with smaller soma size have high-frequency activity, suggesting a mode of birth order dependent microcircuits. Nicotine increased activity frequency and *cholineacetyltransferase-a* (*chata*) mutants showed increased variability in neuronal activity, supporting a role for cholinergic signaling in maintaining stable rhythmic activity. Overall, we reveal the emergence of birth order specified functional microcircuits and show a conserved role of cholinergic signaling in promoting rhythmic neuronal activity in the embryonic vertebrate motor circuit during SNA.

## Materials and Methods

### Zebrafishg

Adult fish were reared at 28°C in a 14/10h light/dark cycle in a fish facility approved by the French Service for animal protection and health (A-75-05-25), and the experiments were performed in compliance with relevant regulations and guidelines. Fish were paired overnight using separators and dividers in mating tanks, and the eggs were collected the following morning after removing the separators. Wild-type AB strain, the *cholineacetyltransferase-a* (*chata*) mutant fish (Granato et al., 1996), *Tg(mnx1:GFP)* (Flanagan-Steet et al., 2005), *Tg(NeuroD:GCaMP6f^icm05^)* (Rupprecht et al., 2016), *Tg(elavl3:H2B-GCaMP6f)* (Chen et al., 2013), *Tg(elavl3:GCaMP6s^CY14^) Tg(elavl3:GCaMP6s^CY44^)* (Vladimirov et al., 2014), *Tg(elavl3:jRGECO1b)* (Mu et al., 2019)*, Tg(elavl3:GCaMP6f)^a12200^* (Wolf et al., 2017), *Tg(mnx1:Gal4)* (Flanagan-Steet et al., 2005), *Tg(UAS :RFP)* (Asakawa & Kawakami, 2009) were used in this study.

### Fluorescent in situ hybridization with antibody labeling

Fluorescent in situ hybridization (FISH) was performed on 24 hpf embryos as described in (Rima et al., 2020). Not1 and Sp6 were used to linearize and synthesize the UTP-digoxigenin labelled *chrna7* probe, respectively. Briefly, dehydrated embryos in methanol were rehydrated by successive washes of decreasing ratio of methanol:PBS solutions. They were then permeabilized and incubated in 50% formamide containing hybridization mixture for three hours before an overnight incubation with UTP-digoxigenin labelled probes. Probes were then detected using horseradish peroxidase conjugated anti-Digoxigenin antibody (Roche) in Maleic acid blocking buffer (150 mM Maleic acid, pH 7.5 & 100 mM NaCl). Signal amplification was performed using homemade tyramide-FITC as described in (Davidson & Keller, 1999). Embryos were incubated with tyramide-FITC (1:250) in TSA reaction buffer (100 mM borate pH 8.5, 0.1% tween-20, 2% dextran sulfate, 0.003% H2O2, and 200 µg/ml 4-iodophenol) for 45 min in dark. The reaction was stopped by washing in 0.1% Tween-20 in PBS (PBT). Embryos were incubated overnight at 4 °C in PBS, 0.1% Triton, 10% sheep serum with rabbit anti-GFP antibody (Thermofisher) to detect GFP. Embryos were washed in PBT and incubated at 4 °C overnight with goat anti-rabbit Alexa594 antibody (Thermofisher). Finally, embryos were washed and mounted laterally in a drop of low melt agarose (2%) and imaged using the Leica SP5 confocal microscope using a 20X water immersion objective.

### Genotyping

DNA was extracted from tail biopsies (adults) or whole embryos using DNA extraction buffer (10 mM Tris pH 8.2, 10 mM EDTA, 200 mM NaCl, 0.5% SDS, 200 µg/ml proteinase K). PCR amplification of the *chata* gene was carried out using primers from (Wang et al., 2008) and the mutation was confirmed by sequencing.

### Pharmacology

Nicotine (Acros Organics) was prepared at a concentration of 10 µM in E3 the day of the experiment. E3 was perfused into the petri dish during the first 15 minutes of acquisition, followed by nicotine for the next 15 minutes using a perfusion peristaltic pump (Ismatec) at a rate of 1.47 mL/min. The tubes of the peristaltic pump were thoroughly cleaned with E3 between each acquisition.

### Calcium imaging

Embryos were individually transferred to a 60 mm petri dish and mounted using a drop of TopVision Low Melting Point Agarose (2% in E3, ThermoFisher) with pancuronium bromide (300 µM). Calcium imaging recordings were carried out on a spinning disk microscope (Zeiss Axio Examiner.Z1) using a 40X water immersion objective (NA = 0.95) acquired at a rate of 3.3 Hz [*Tg(elavl3:GCaMP*)] or 10 Hz [*Tg(elavl3:GCaMP;mnx1:gal4;UAS:RFP*)] using the 480 nm laser. The 532 nm laser was used for the z-stack acquisition of mnx1:RFP^+^ cells. The duration of acquisition ranged from 12 – 30 min depending on the experiment.

### Calcium imaging analysis

#### FIJI

Regions of interests (ROIs) were manually drawn on the soma of neurons exhibiting GCaMP fluorescent transients and the average fluorescence intensity was extracted for all recorded frames using FIJI (Schindelin et al., 2012). Further analysis was done on MATLAB (The MathWorks, Inc. 2022).

#### MATLAB

The values for the average fluorescence intensity for each ROI per frame were imported into MATLAB using custom written scripts. Z-score was used to analyze calcium activity according to the formula:

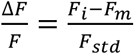

*F*_*i*_ is the mean intensity in a single ROI at a single time point while *F*_*m*_ is the moving mean in a single ROI for 500 frames and *F_std_* is the standard deviation for the entire recording. Clustering of neurons was performed using *k-means* function and manually verified based on pairwise linear correlation coefficients. Each calcium event was identified using *findpeaks* (Signal Processing Toolbox, MATLAB) function in MATLAB and verified manually. All plots and graphs were made on MATLAB.

Alternation index was calculated as:

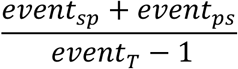

where *event*_*sp*_ is the total number of time when calcium transient in the ipsilateral soma is followed by an event in the puncta, *event_ps_* is the total number of time when calcium transient in the puncta is followed by an event in the ipsilateral soma and *event_T_* is the sum of events in the ipsilateral soma and puncta.

Symmetry index was calculated as:

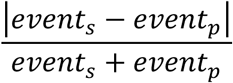

where *event_s_* is the total number of events in the ipsilateral soma, *event_p_* being the total number of events in the puncta displaying contralateral activity.

### Motoneuron soma size analysis

The calcium activity of each elavl3:GCaMP;mnx1:RFP^+^ cells was first categorized into low frequency (LF), high frequency (HF) or no activity (NA). Each neuron soma was outlined manually on a Z-stack image of the mnx1:RFP^+^ cells and its fluorescence intensity and area quantified in FIJI. Fluorescence intensity was normalized by dividing the values by that of the largest CaP neuron per embryo.

### Muscle contraction, somite imaging and analysis

Microscope and mounting setup were identical to calcium imaging (10 Hz) with the exception of using transmitted light to image muscle contractions in non-paralyzed embryos mounted in low melting point agarose (2% in E3). Prior to imaging muscle contractions, z-stack acquisition of mnx1:RFP^+^ cells was performed using the 532 nm laser. The recordings were analyzed using FIJI where a ROI was drawn in an area that displayed a distinct morphology (*i.e*., muscle segment, edge of embryo, etc.) and the change in pixel intensity upon movement was quantified. Z-stack images were acquired using transmitted light and 532 nm laser at different anterior-posterior axis of the embryo and the number of somites and mnx1:RFP^+^ cells were manually counted.

### Coiling behavior, heartbeat, length measurements and analysis

Coiling behavior of dechorionated embryos was recorded at 25 Hz using a color camera (Leica IC80 HD) mounted on a stereomicroscope (Leica M80). For measurements of *chata* mutant and WT siblings, embryos from a cross between *chata^+/-^* fish were phenotyped into moving (WT sibling) or non-moving (mutant). To record the heartbeats, the embryos were laterally mounted in 1.5% low melt agarose. The number of heartbeats was manually counted for the duration of the 1 min recording. To analyze the length of the embryos, images were acquired of embryos paralyzed using tricaine. To calculate the length of the embryos, the outline of the embryo from the nose tip to the tail tip were traced using FIJI.

### Statistics

Statistical comparisons between two independent groups were conducted using a Wilcoxon rank sum test or a Student’s t-test. Comparison between more than 2 dependent groups were conducted by a Kruskal-Wallis test followed by a Bonferroni test. Statistical tests were performed on MATLAB. Specific information on statistical test, sample size, value representation (mean ± std) and p values are indicated in the figure legends.

## Results

### Spinal neurons display different patterns of spontaneous activity

To better understand the mechano-dynamics of the left-right coiling behavior, we first analyzed the motor behavior of dechorionated embryos. We found that the first visually identifiable movements occurred in the head and tail areas. The tail continued to contract, which was then followed by slow extension of the tail (Figure 1). The area between the caudal head and rostral trunk showed the least visible movement during the coiling behavior, suggesting that the bending of the body pivots around this region. As this was the most stable region along the body axis, we decided to analyze the neuronal activity in this region, which we will refer to as the pivot point.

**Figure 1.**
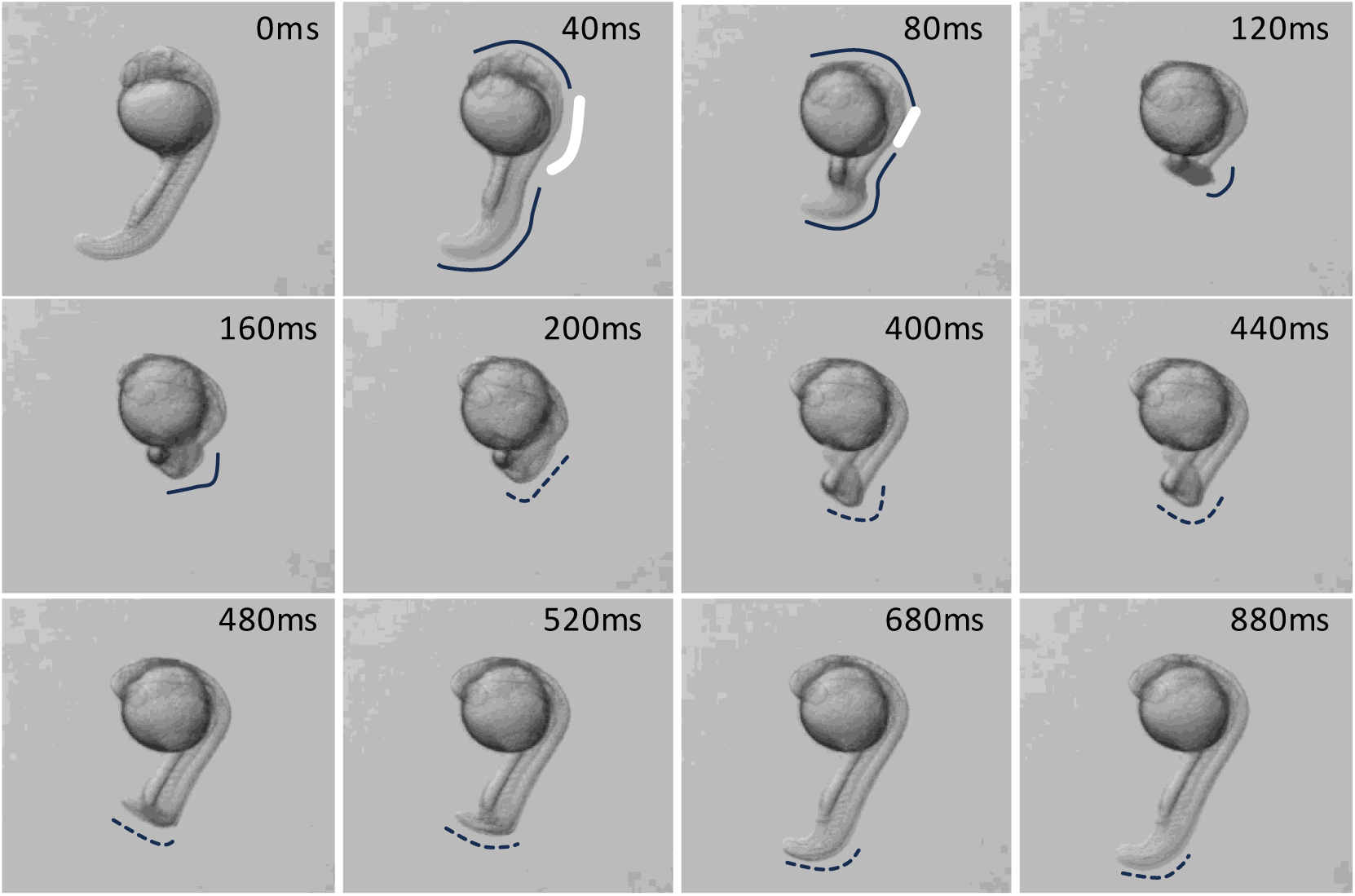
1-day old embryos show distinct movements around the pivot point during coiling behavior. A representative time lapse recording of a 1-day old embryo showing specific movements along the rostral-caudal body axis. Movements along the body axis are easily detectable in the head and the tail areas while the trunk (pivot point, white line) shows less movement. Location of movements are outlined along the body axis depicting muscle contractions (solid line) and relaxations (dotted line). N = 16 embryos, n = 32 single coiling events.

We first carried out calcium imaging on embryos expressing GCaMP under different pan-neuronal promoters. While all 6 transgenic lines displayed calcium transients at 5 days, only one line [*Tg(elavl3:GCaMP6f ^12200a^)*] (Wolf et al., 2017) showed activity in 1-day old embryos (Table S1). The result was surprising as at least two other transgenic lines were shown to display calcium activity in 1-day old embryos in a previous study (Wan et al., 2019). The results suggest that the signal-to-noise ratio of GCaMP activity differ between individuals but becomes more robust during development.

We carried out calcium imaging of the pivot point in the rostral spinal cord (above the beginning of the yolk sac extension) in 1-day old laterally mounted *Tg(elavl3:GCaMP6f ^12200a^)* embryos expressing the GCaMP transgene in most spinal neurons, including the motoneurons (Figure 2A). We found that around 35 % (34.5 ± 8.9) of imaged neurons showed at least one calcium event during the 10 min recording (Figure 2B). We analyzed activity in the ventral domain containing motoneuron and interneurons vs. dorsal domain consisting mainly of sensory neurons. Sensory neurons are easily identifiable due to their rounder soma (Williams & Ribera, 2020) and expression of higher baseline GCaMP fluorescence. We found that the ventral spinal cord contained more active neurons compared with the dorsal domain (Figure 2A, B), consistent with what has been reported in other studies (Saint-Amant & Drapeau, 2001). However, in contrast to previous studies showing that all active spinal neurons display synchronized activity (Saint-Amant & Drapeau, 2001; Wan et al., 2019; Warp et al., 2012), we observed groups of neurons exhibiting distinct patterns of activity. We carried out *k-means* clustering analysis and determined that *k* = 3 was sufficient to group neurons into highly correlated clusters with each cluster containing at least 10% of active neurons (Figure 2C-E). Neurons in the first two clusters showed highly correlated activity within the cluster, while they were less correlated in the third cluster (Figure 2D, E). We next analyzed the location of the neurons and found that largely neurons in the first and second groups were intermingled in the ventral spinal cord, whereas neurons in the third group were located mostly in the dorsal area (Figure 2C). As motoneurons are largely restricted to the ventral-most spinal cord, we hypothesize that neurons in the first and second clusters are composed of either interneurons or motoneurons or a combination of both types.

**Figure 2.**
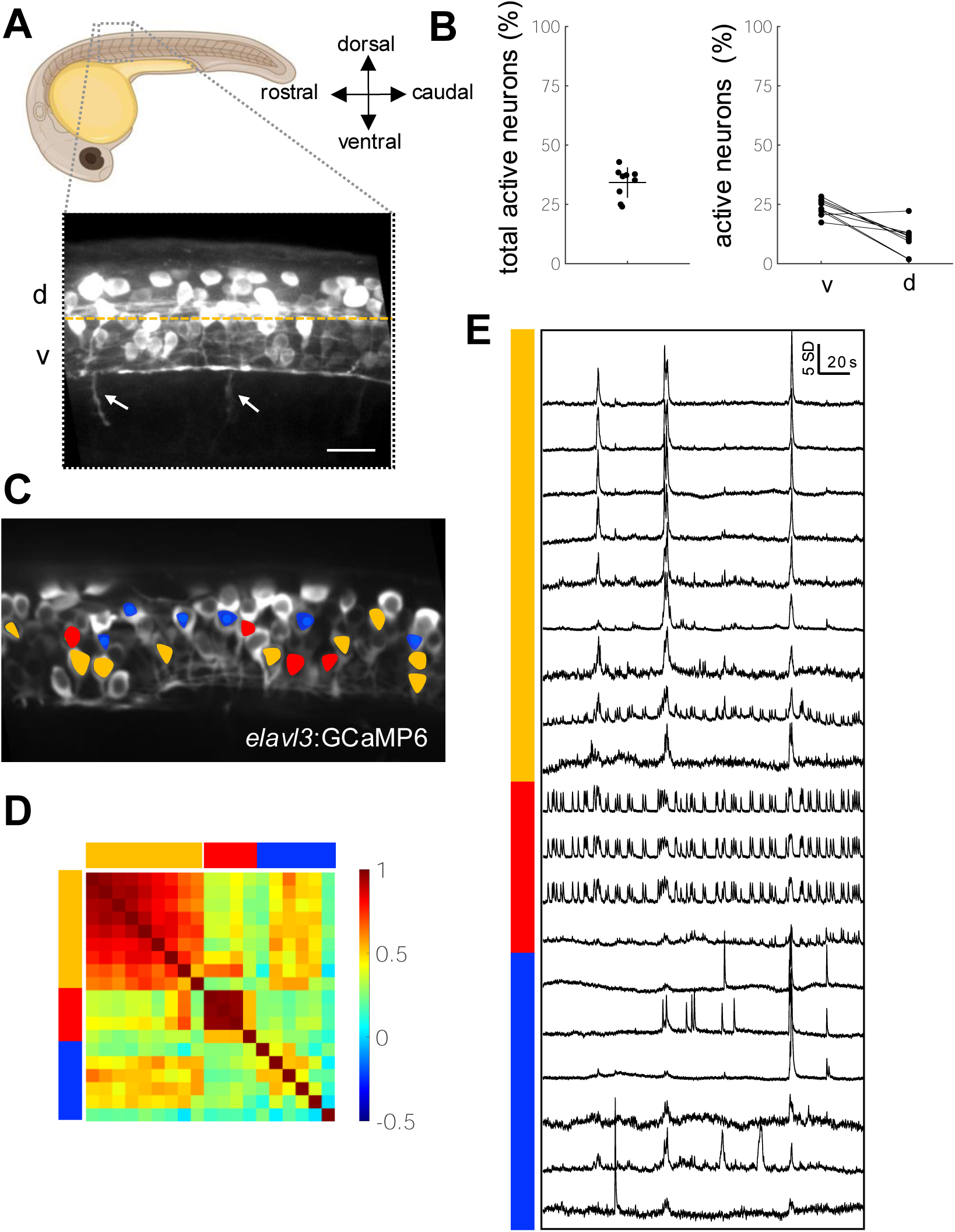
1-day old embryos display different patterns of spontaneous activity. (A) Schematic of a lateral view of a 1-day old embryo indicating the calcium imaging area (dotted box). Close up z composite image of the recorded area along the spinal cord in a *Tg(elavl3:GCaMP6*) embryo. Orange horizontal line demarcates dorsal (d) vs. ventral (v) regions quantified in (B). Note the motoneuron axons exiting the spinal cord (arrows). Double arrowheads indicate rostral, caudal, dorsal, ventral axis of the embryo. Scale bar = 20 µm. (B) Graphs showing the percentage of total *elavl3:GCaMP^+^* cells displaying calcium events (active neurons) 34.2 ± 6.37% (left) and those located in the dorsal (23.6 ± 3.4%) vs. ventral (10.6 ± 6.1%) spinal cord domains (right). mean ± SD. N = 9 embryos, n_T_ = 419 total neurons, n_A_ = 144 active neurons. (C-E) *k-*means clustering (*k* = 3) reveals neuron groups that display distinct patterns of calcium activity in a representative 1 day old *Tg(elavl3:GCaMP6*) embryo. 8 min recording at 3Hz. (C) Spinning disk image of the imaged region in a representative embryo showing the location of the different neuron groups overlaid by colored circles. (D) Correlation coefficient matrix of active neurons indicating the degree of correlation between neurons within each group and with other groups. (E) Z-score plot of the 3 groups of neurons revealed by *k*-means clustering. Neurons in different clusters are indicated by the color-coded bar on left.

### Circuits with high or low frequency activity are recruited during spontaneous activity

To determine the cell types in the first two neuronal clusters, we recorded calcium activity in embryos that co-express red fluorescent protein (RFP) mainly in motoneurons [*Tg(elavl3:GCaMPf;mnx1:gal4;UAS:RFP*)] (Asakawa & Kawakami, 2009; Flanagan-Steet et al., 2005). The three motoneurons that are born first in each hemisegment of the spinal cord are the primary motoneurons, CaP, MiP and RoP, that bundle their axons and exit the spinal cord together to target the ventral, dorsal and medial muscle fibers, respectively (Figure 3A) (Moreno & Ribera, 2009; Myers, 1985). Next, secondary motoneurons are generated between the primary motoneurons and display smaller soma size (Myers et al., 1986). Zebrafish embryos undergo rapid development, with approximately 2 somites added to their trunk every hour during somitogenesis (D’Costa & Shepherd, 2009). To determine the developmental stage of the embryos, we quantified the relationship between the number of somites and the number of mnx:RFP+ cells per motoneuron bundles in the pivot point (Figure S1). As expected, we found that the number of mnx:RFP+ cells increased with somite number. We found that embryos with 20-25 somites (19-21 hours post fertilization (hpf)) contain 1-3 RFP^+^ cells per motoneuron bundle, suggesting that the RFP^+^ cells are primary motoneurons (PM). 26-30 somite stage embryos (22-24 hpf) contain 4-7 RFP^+^ cells, containing both primary and secondary motoneurons (PSM). We will henceforth refer to the embryos with RFP^+^ primary motoneurons as PM embryos and those expressing RFP in both primary and secondary motoneurons as PSM embryos.

**Figure 3.**
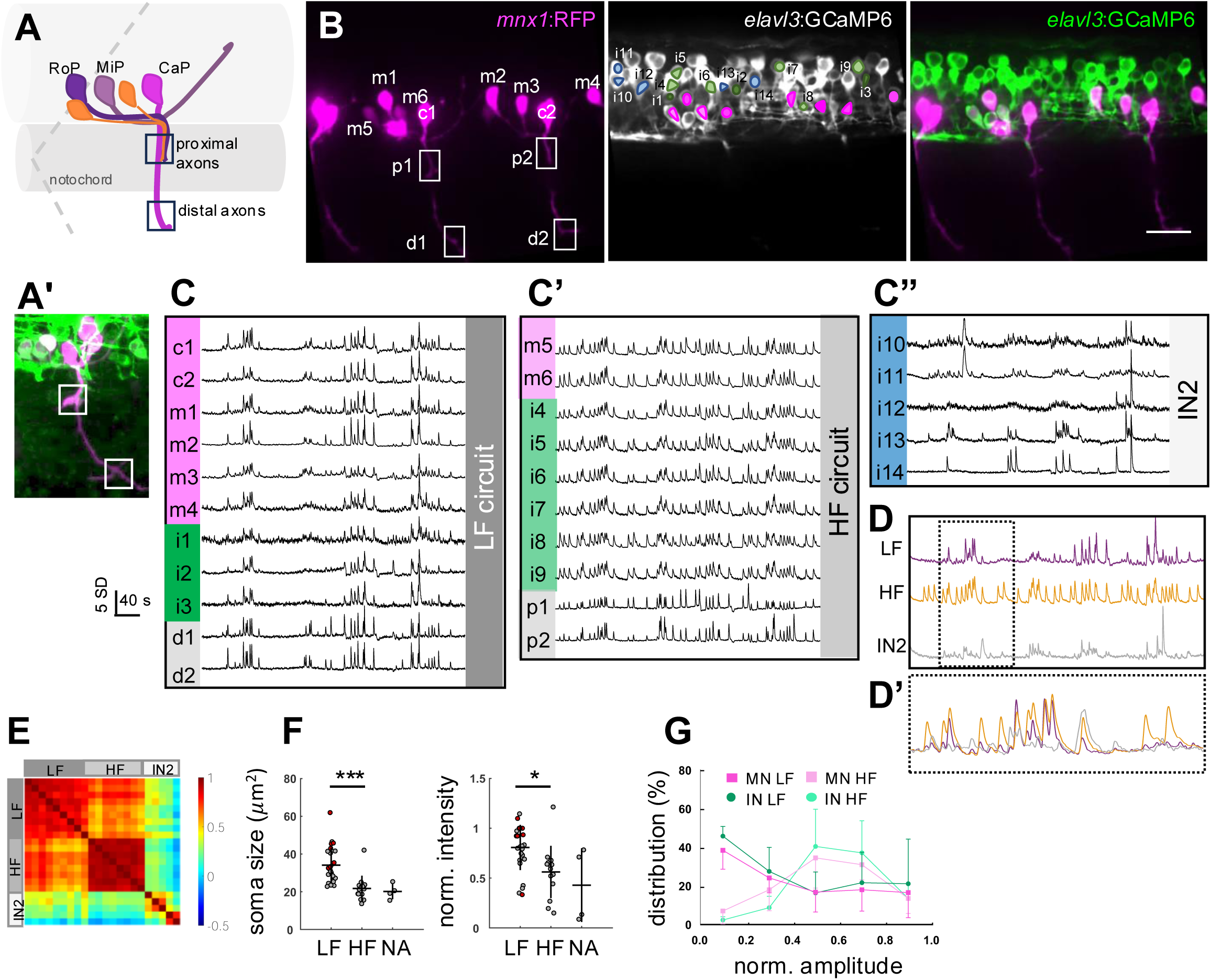
Functionally distinct microcircuits are present during spontaneous activity. (A,A’) Schematic of primary (CaP, MiP, RoP) and secondary (orange) motoneurons and their axon projections displaying stereotypical organization (A). Note that the proximal axons represent those from both the primary and secondary motoneurons, whereas, the distal axons represent mainly the CaP axon in PSM embryos. p: proximal, d: distal axons. (B) Spinning disk image of the recorded area in the spinal cord of a representative *Tg(elavl3:GCaMP6 ; mnx1:RFP)* embryo showing *mnx1:RFP*^+^ (left panel, magenta), *elavl3:GCaMP6^+^* (middle panel, white) and *elavl3:GCaMP6 ; mnx1:RFP*^+^ (right panel) neurons. To visualize the axons, the image for *mnx1:RFP*^+^ labeling is a composite image of 7.5 µm in the z axis. Neuron type is color-coded (motoneurons: magenta, interneurons: green, blue) and identifiers noted (c: CaP, m: motoneuron, i: interneuron, p: proximal axons, d: distal axons). Scale bar = 20 µm. (C-C”) Activity plot of a 5-minute calcium imaging recording. Neurons types are color-coded (motoneurons: magenta, interneurons: green, blue, axons: grey) and identifiers noted to the left of the panel (m: motoneuron, c: CaP, i: interneuron, p: proximal axons, d: distal axons). Neurons belonging to distinct circuits are color-coded in different shades of grey to the right of the panel (HF: high frequency circuit, LF: low frequency circuit). (D,D’) Average plots of neurons in LF (purple), HF (orange) circuits and IN2 type (grey) (D), and close up overlay of the average plots in dotted rectangle (D’). (E) Pairwise correlation coefficient matrix between neurons. Neurons in functional circuits are indicated by grey bars left and above the matrix. (F) Scatter plot of soma size (left) and fluorescence intensity (right) of *mnx1:RFP*^+^ neurons showing LF, HF or NA. CaP neurons are indicated in red. N = 6 embryos, n = 26 LF, n = 14 HF, and n = 4 non-active neurons. Soma size : LF = 34 ± 9.4 µm^2^, HF = 21.7 ± 6.7 µm^2^. Normalized intensity : LF = 0.81 ± 0.23, HF = 0.56 ± 0.26. Mean ± 0.37 SD. Student’s t test. * <0.05, ***<0.0001. (G) Histogram showing the distribution of normalized amplitudes from MNs and INs in the HF and LF circuits. N=10 embryos, n = 1101 events. Also see Figure S1.

Consistent with our previous observation (Figure 2), we found that *k*-means clustering analysis separated the spinal neurons into 3 different clusters based on their activity pattern in PSM embryos with 4-5 motoneurons per bundle (Figure 3B-E). The first cluster of neurons displayed low frequency (LF) activity whereas the second cluster of neurons showed high frequency (HF) activity (Figure 3C-C”). The two clusters contained both RFP^+^ and RFP^-^ neurons (Figure 3B-C”), confirming the presence of distinct microcircuits containing motoneurons and interneurons that display either LF or HF activity (Video S1). The third neuronal cluster showed low correlation activity with the first two clusters and did not contain any RFP^+^ cells (Figure 3B, E), suggesting that these are a different interneuron type (IN2) that are not integrated into the LF or HF circuits.

One explanation for the presence of distinct functional microcircuits is that they consist of neurons that are born and recruited at different times. First-born primary CaP motoneurons are easily identified based on the location of the soma directly over the axon bundle exiting the spinal cord. We found that the RFP^+^ neurons that showed such morphological criteria (Figure 3B: c1, c2) consistently displayed LF activity (Figure 3A-C). This was also the case with other neurons that had larger soma size and strong RFP expression (Figure 3B, C: m1, m2, m3). By contrast, neurons with smaller soma and faint RFP expression showed HF activity (Figure 3A-C’: m4, m5). Quantification of size and intensity of the soma in RFP^+^ cells show a clear difference between those with HF vs. LF activity.

The distal part of the motoneuron axon bundles contains mainly CaP axons whereas the proximal motoneuron axon bundles contain MiP and RoP as well as later-born secondary motoneuron axons (Figure 3A, A’). If the LF circuit consists of earlier born neurons while the HF circuit contains later born neurons, we expected to observe LF activity in the distal axons and HF activity in the proximal axons. Indeed, we found that axons in the distal area showed LF activity while those in the proximal area displayed HF activity (Figure 3B-C’), suggesting that the LF and HF circuits consist of earlier born (older) and later born (younger) motoneurons, respectively.

Overlaying the average traces of calcium activity between the LF and HF circuits revealed that when neurons displaying LF activity are active, neurons in the HF circuit are also active. However, calcium events that occur in neurons with HF activity are not always observed in neurons with LF activity (Figure 3D, D’). We next analyzed the amplitude of calcium events in PSM embryos and found that the HF circuit consist of events that were more uniform between each other, whereas neuron activity in the LF circuit showed events with variable amplitudes (Figure 3F). Taken together, we hypothesize that the LF and HF circuits are not independent from one another but rather that the LF activity emerges from HF activity by gradual uncoupling of events.

### Early spontaneous activity consists of synchronized high frequency rhythmic activity

If PSM embryos contain a LF microcircuit composed of early-born older neurons and a HF microcircuit composed of late-born younger neurons, we expected that PM embryos would only have one circuit exhibiting HF activity. We analyzed the frequency and pattern of calcium activity in PM embryos. Indeed, we found that the mnx:RFP^+^ motoneurons only displayed one type of calcium activity that showed HF rhythmic activity (Figure 4A-D). We found two types of interneuron activity consisting of one that synchronized (IN1) and another that had no significant correlation (IN2) with motoneuron activity (Figure 4A-D).

**Figure 4.**
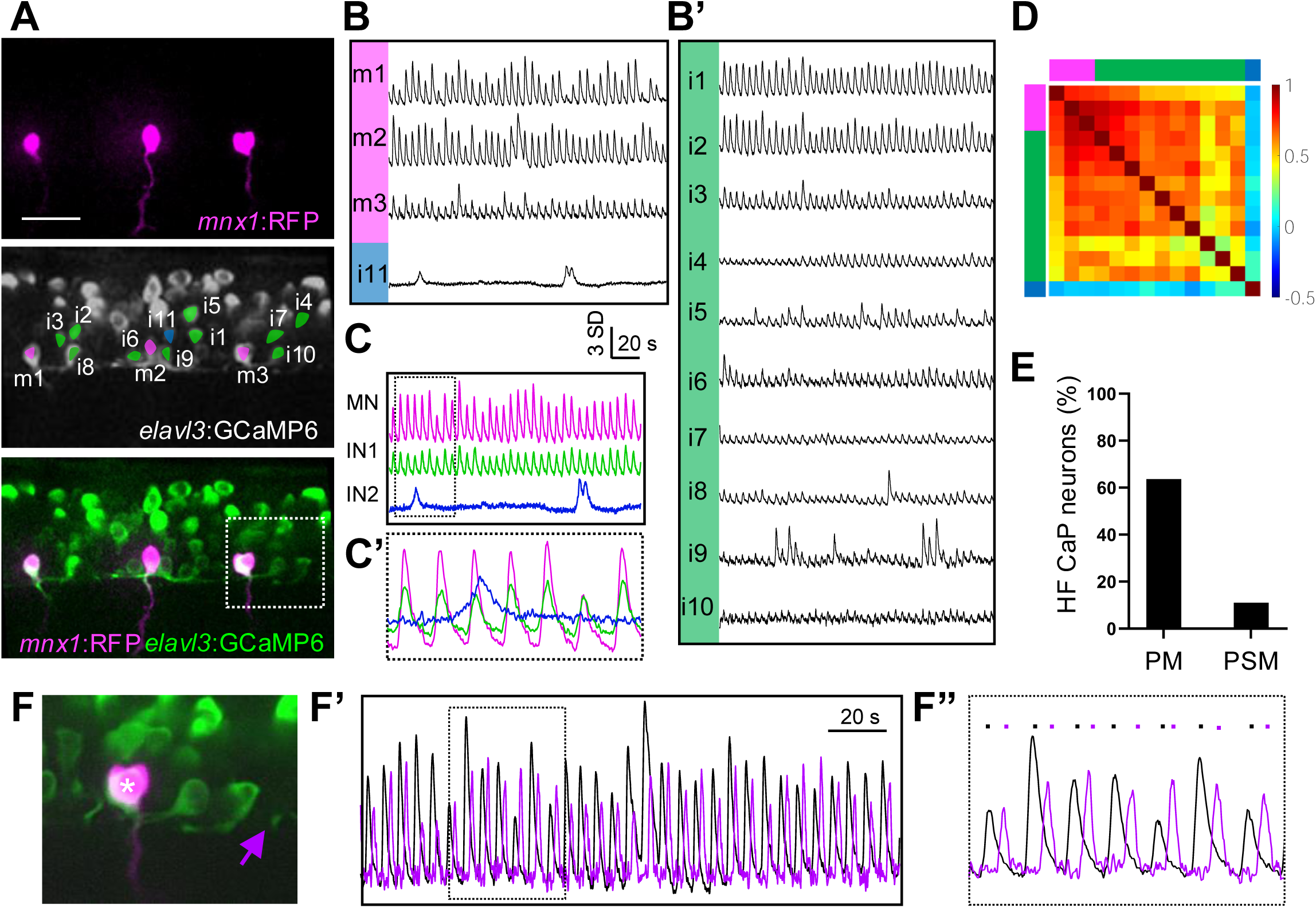
Early-stage embryos exhibit fast rhythmic activity. (A) Spinning disk image of the recorded area in the spinal cord of a representative *Tg(elavl3:GCaMP6 ; mnx1:RFP)* embryo showing *mnx1:RFP^+^* (magenta, top panel), *elavl3:GCaMP6^+^*(white, middle panel) and *elavl3:GCaMP6 ; mnx1:RFP* ^+^ (bottom panel) neurons. To visualize the axons, the image for *mnx1:RFP*^+^ labeling is a composite image of 7.5 µm in the z axis. Active neuron soma of motoneurons (MNs, magenta), interneuron type 1 (IN1, green), interneuron type 2 (IN2, blue) and their identifiers are indicated (middle panel). Scale bar = 20 µm. (B,B’) Z-score plot of a 3.5 minute calcium activity recording. Neurons are ordered by type: MNs (magenta), interneurons that display correlated activity with motoneurons (IN1, green) and interneuron with no correlation to motoneuron activity (IN2); blue). Neuron type (colored bars on left) and identifiers are indicated. (C,C’) Average (D) and close up overlay (D’, dotted rectangle) activity plots of MNs (magenta), IN1 (green) and IN2 (blue). (D) Correlation coefficient matrix of active neurons. MNs (magenta), IN1 (green), and IN2 (blue) are are indicated by colored bars on top and left of the matrix. (E) Bar graph displaying the percentage of primary CaP neurons displaying HF activity in early vs late embryos. PM embryos: N=7, n = 11 CaP neurons ; PSM embryos: N = 10, n = 17 CaP neurons. (F-F”) (F) Close up image of MN soma (asterisk) and puncta (purple arrow) outlined in (A) that display alternating calcium activity. MN soma (black plot) and puncta (purple plot) (F’). (F”) Close up of activity plot in the dotted rectangle in (F’). Colored circles above the activity plot indicate soma (black) or puncta (purple) activity.

Since CaP motoneurons are the first-born easily identifiable neurons in the spinal cord, we next analyzed the activity type in the PSM embryos. We expected that in younger embryos, the CaP neurons would show HF activity, whereas in older embryos would show LF activity. Indeed, we found that CaP neurons in younger PM embryos exhibit largely HF activity, whereas older PSM embryos mainly showed LF activity (Figure 4E). The result strongly suggests that newly-born younger neurons have HF activity that transitions to LF activity as the neurons mature. One noteworthy observation was the presence of puncta and/or axons that showed activity that alternated with that in the neuron soma in the rhythmic HF circuit (Figure 4F-F”, Video S2). One likely explanation for the alternating activity in these puncta and/or axons is that they are due to the activation of the contralateral motor circuit.

### Spontaneous calcium activity corresponds to trunk contractions

To address whether the alternating activity in the puncta/axons corresponds to contralateral muscle contractions, we first analyzed muscle contractions in non-paralyzed agarose-restrained embryos expressing RFP in motoneurons. In PM embryos, we identified two types of muscle contractions that showed a large vs. small movement. The two movements alternated around every 2 sec (0.5Hz), suggesting that they correspond to the ipsilateral vs. contralateral muscle contractions that occur during left-right coiling (Figure 5A-B). We quantified the relationship between the frequency of muscle contractions with the number of motoneurons per bundle and found they have an inverse correlation (Figure 5E), similar to what was observed during spontaneous neural activity in spinal neurons (Saint-Amant & Drapeau, 1998). Next, to confirm that the calcium activity in motoneurons correspond to muscle contractions, we carried out simultaneous calcium imaging of motoneurons with muscle contractions (Figure 5C-D). We found that calcium activity in motoneurons co-occurred with around half of the muscle contractions. The result suggests that the motoneuron activity induces ipsilateral muscle contractions while the muscle contractions that occur without calcium activity are due to the contralateral motoneuron activity. Altogether, the results demonstrate that the calcium activity in the motor circuit translates to muscle contractions and suggest that the alternating calcium activity observed in the puncta and/or axons corresponds to the activation of the contralateral motor circuit.

**Figure 5.**
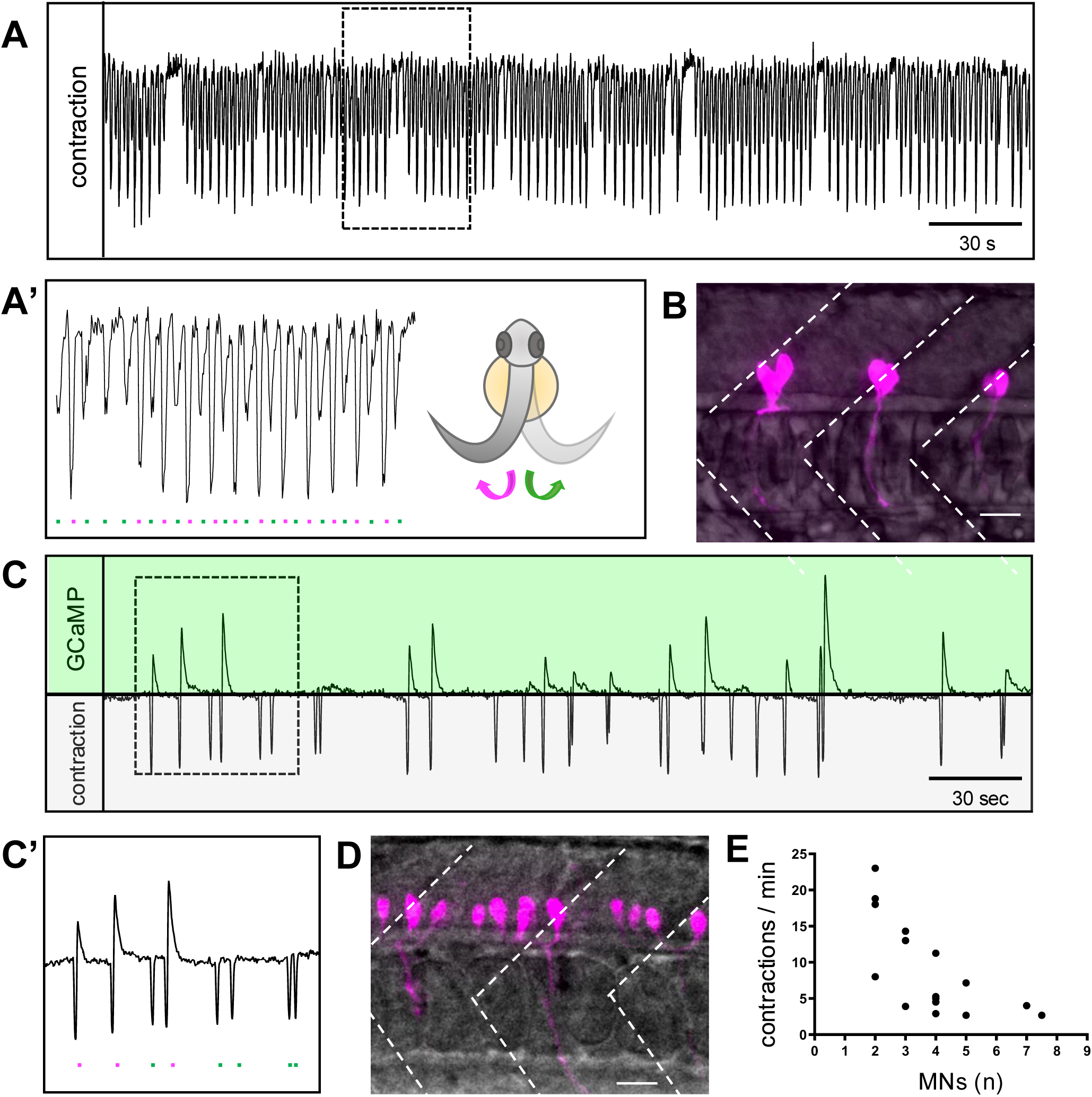
Muscle contractions correlate with motoneuron calcium activity and frequency. Muscle contractions during a 5-minute recording of an early-stage embryo. (A’) Zoomed in muscle activity plot from time shown in (A, rectangle). Putative ipsilateral (magenta) and contralateral (green) contractions are indicated by colored circles. Cartoon of zebrafish embryo from a frontal view showing left-right coiling corresponding to ipsilateral (magenta) or contralateral (green) contractions. (B) Lateral view showing motoneurons (magenta) and muscle (grey) in an early stage *Tg(mnx1:RFP)* embryo containing 2 MNs per hemisegment. White dotted lines delineate the somite boundaries. (C) Ipsilateral motoneuron activity (top, green) and muscle contraction (bottom, grey) during a 5 minute recording of a late-stage embryo. Scale bar = 20 µm. (C’) Zoomed in muscle activity plot from time shown in (D, rectangle). Ipsilateral (magenta) and contralateral (green) contractions are indicated by colored circles. (D) Lateral view showing motoneurons (magenta) and muscle (grey) in late-stage embryo containing 5 MNs per hemisegment. White dotted lines delineate the somite boundaries. Scale bar = 20 µm. (E) Scatter plot showing negative correlation between developmental stage (identified by number of *mnx1^+^*cells per hemisegment) and frequency of muscle contraction. N=15 embryos.

### Nicotinic receptors modulate high frequency activity

Nicotinic acetylcholine receptor (nAChR) subunits α2 and α3 are expressed in the INs and α7 in motoneurons as early as 20 hpf (Rima et al., 2020). We sought to determine whether cholinergic signaling, via the nAChRs, could affect early neuronal activity. We carried out calcium imaging while perfusing the embryo with nicotine or control medium (E3). In PSM embryos, spinal neurons did not show any changes in their activity patterns during the first 5 min following nicotine perfusion. However, between 5-10 min in nicotine, the frequency of calcium events increased by around 1.3-fold in both HF and LF circuits. After 10-15 min in nicotine, some embryos showed further increased activity whereas others became inhibited (Figure 6A-B’). The results show that activating nAChRs via nicotine in PSM embryos increase their activity, suggesting a role for cholinergic signaling during SNA.

**Figure 6.**
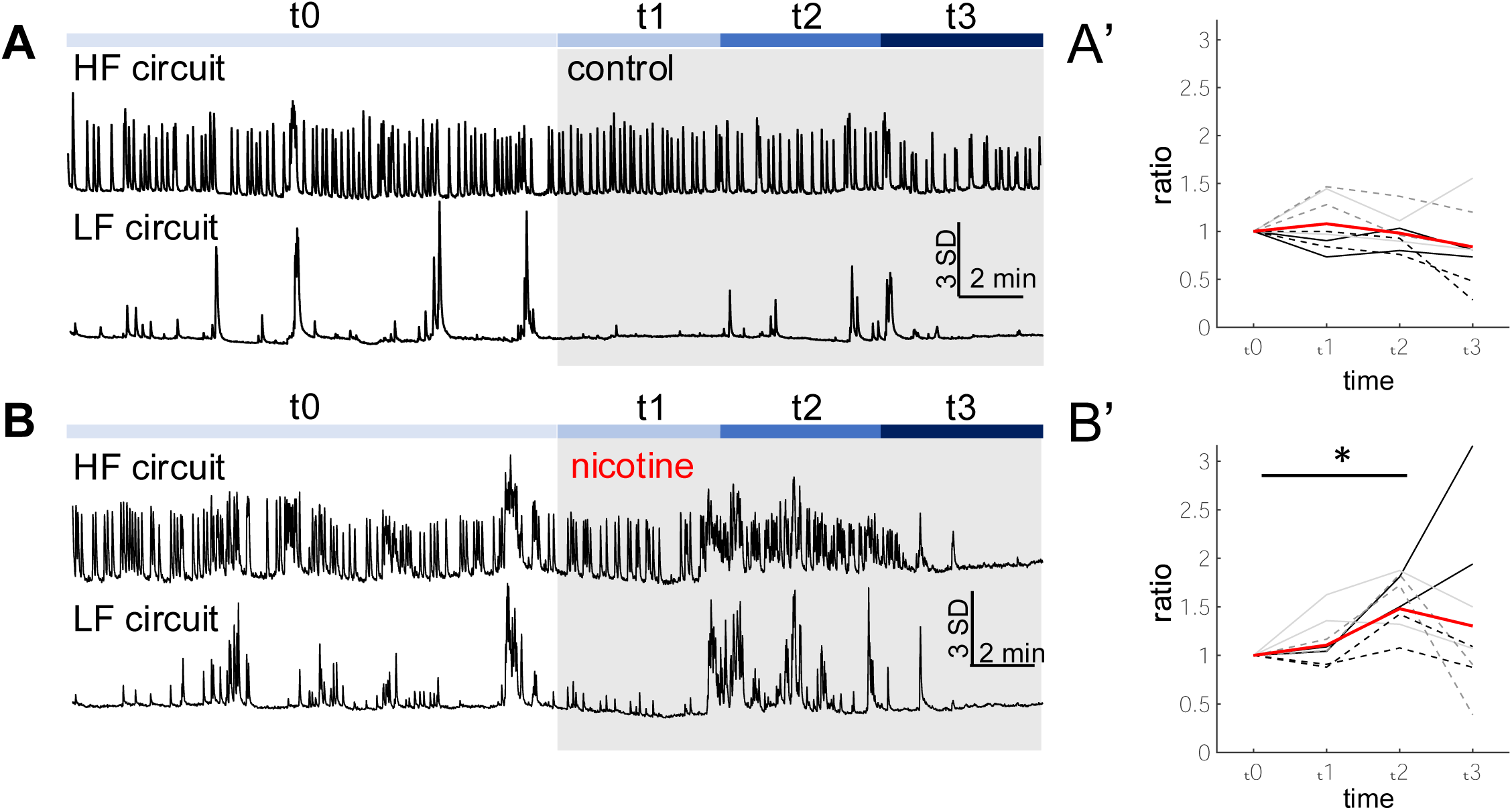
Nicotine increases spinal neuron activity during SNA. (A, B) Activity plots of representative neurons in the high frequency (HF) and low frequency (LF) circuits during baseline (t0), 0-5 min (t1), 5-10 min (t2), and 10-15 min (t3) after the start of perfusing control E3 medium (A) or nicotine (B). (A’, B’) Graph showing the number of calcium events normalized by the number of events during t0 for the four sections (t0, t1, t2, t3, t4). Each color and line type represent events in HF and LF circuits in the same embryo. Mean values are plotted in red. (B’) t2/t0 = 1.48 ± 0.32 (mean ± std). N = 4 (control), N = 5 (nicotine) HF and LF circuits analyzed for each embryo. Kruskal-Wallis followed by Bonferroni test * p = 0.0055.

### Cholinergic signaling is required for alternating rhythmic activity

*Cholineacetyltransferase-a* (*chata*), which encodes an enzyme for acetylcholine synthesis, is expressed in motoneurons in 1-day old embryos (Rima et al., 2020). We next investigated whether cholinergic signaling plays a role during SNA by carrying out calcium imaging in *chata* mutants (Wang et al., 2008) expressing GCaMP in most neurons and RFP in motoneurons [*Tg(elavl3:GCaMP6f;mnx1:gal4;UAS:RFP*); *chata^-/-^*]. First, we confirmed that there was no overt difference in the development of the embryos by measuring the length and heartbeat frequency between the mutant and WT siblings (Figure S2A, B). Next, we quantified the percentage of active cells and found no significant difference between mutant and WT siblings (29.2 ± 16.7% vs. 34.5 ± 7.9 %) (Figure S2C). We analyzed the calcium activity pattern and also found similar percentages of active motoneurons, IN1 and IN2 in the *chata* mutants compared to their WT siblings (Figure S2D). We found that with the exception of one PM embryo, *chata* mutant embryos showed one type of HF rhythmic activity, similar to that observed in WT siblings. In older PSM embryos both *chata* mutant and WT siblings displayed distinct functional circuits with HF and LF activity (Figure S2E).

However, careful inspection of the calcium events revealed that the HF rhythmic activity in PM embryos appeared less regular in *chata* mutants compared to WT siblings (Figure 7A-A’). Indeed, analysis of the interval time between consecutive calcium events in the *chata* mutant neurons revealed a higher degree of variability compared to that in the WT neurons (Figure 7A-B). We wondered whether the longer interval time between events was due to the absence of general activity or rather because the contralateral side was active during this time. We analyzed calcium activity in the soma vs. puncta to infer the activation of ipsilateral vs. contralateral motor circuits. The alternation index was calculated so that if there is perfect alternation between the ipsi- and contralateral sides, the alternation index would be 1. In WT embryos, we were able to fit a linear regression model between the alternation index and number of motoneurons, showing a decrease in the frequency of left-right coiling during development (Figure 7C-D). By contrast, we found that the *chata* mutants had a lower alternation index compared with WT siblings (Figure 7C-D, Video S3), suggesting that left-right alternating activity is aberrant in the *chata* mutants in PM embryonic stages. Although the left right alternation may be defective in the *chata* mutants, the total number of events in the left vs. right sides could still remain the same. We analyzed the symmetry index where the value is 0 if the number of events between the left and right sides is the same. We found that the *chata* mutants have a higher value for their symmetry index compared with their WT siblings, suggesting that the left and right sides are less symmetrically active in the mutants compared to their WT siblings. These results indicate that cholinergic signaling is necessary for modulating HF rhythmic activity as well as left-right symmetric activity in the zebrafish embryo.

**Figure 7.**
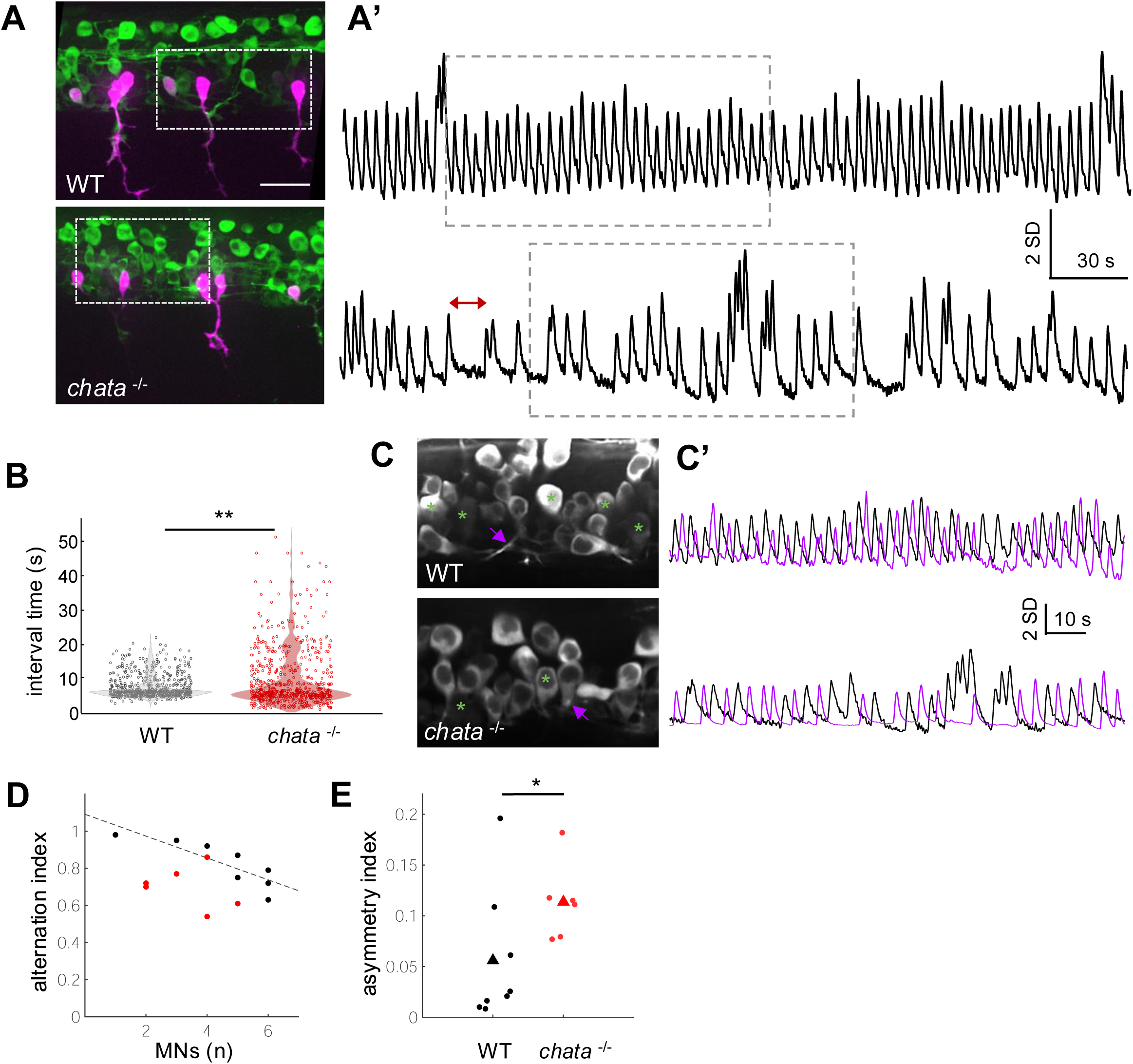
Cholinergic signaling is required for maintaining early rhythmic spontaneous activity. (A,A’) Images of recorded spinal area showing MNs (magenta) (A) with plots of representative neurons in WT (top) or *chata^-/-^* (bottom) embryos (A’). (B) Scatter plots showing the interval time between two consecutive peaks (red double arrow in A’) WT (black) or *chata^-/-^* (red) PM embryos. WT : N = 4 embryos, n = 720 events; *chata^-/-^*: N = 6 embryos, n = 720 events. Wilcoxon rank-sum test ** p = 1.3e-5. (C,C’) Image showing neuron soma (asterisks) and puncta (purple arrow) outlined in white dotted rectangles in (A, A’) that display alternating activity (C’). Close up of calcium activity in soma (black) and puncta (purple) in WT (top) and *chata^-/-^* (bottom) embryos outlined by grey dashed rectangle in A’. (D) Linear regression of alternation index between developing WT (black) and *chat^-/-^* (red) PM and PSM embryos. (E) Asymmetry index show *chata^-/-^* embryos display less left right symmetry in calcium events compared with WT siblings. Each circle represent the asymmetry index for an embryo, triangles show mean value. WT vs. *chata^-/-^* : 0.056 ± 0.062 vs. 0.114 ± 0.038 (mean ± std). WT: N = 8 embryos, n = 555 events, *chat^-/-^* : N = 6 embryos, n = 566 events. Wilcoxon rank-sum test. * p = 0.0256. Also see Figure S2.

## Discussion

### Emergence of distinct functional circuits in the developing embryo

By carrying out high resolution lateral view calcium imaging of spinal neurons, we discovered that distinct IN-MN microcircuits are formed in a birth order dependent manner during coiling behavior. Previous studies have only reported global synchronization of active spinal MN and INs during this critical period (Wan et al., 2019; Warp et al., 2012), suggesting that they focused only on PM embryos with high frequency rhythmic activity.

We found that younger PM embryos displayed one type of HF rhythmic activity, in both mnx1:RFP^+^ and mnx1:RFP-negative neurons, suggesting a single circuit composed of motoneurons and interneurons. Indeed, the result is consistent with previous study showing that CiA, CiD and CoPA interneurons are functionally active and that the CiD interneurons are electrically coupled with CaP motoneurons in 1 day old embryos (Saint-Amant and Drapeau., 2001). Older PSM embryos contained functional motor microcircuits with distinct patterns of low-frequency (LF) or high-frequency (HF) activity. While all events that occur in the HF circuit are not observed in the LF circuit, all activity in the LF circuit co-occurs with those in the HF circuit. In other words, the LF activity is nested within the HF activity. Based on the morphology and topography of motoneuron soma and axons, we found that the older motoneurons display LF activity whereas the younger motoneurons show HF activity. Taken together, we propose a model for the emergence and evolution of spontaneous neural activity (Figure 8). First, upon neurogenesis, a group of primary motoneurons and INs establish a functional circuit consisting of rhythmic HF activity driven by gap junctions, likely coupled to ‘pacemaker cells’, which are thought to be the ipsilaterally caudal projecting (IC) and/or ventrolateral descending (VeLD) cells (Miles et al., 2024; Saint-Amant & Drapeau, 2001; Tong & McDearmid, 2012). As the embryos develop, gap junctions between the neurons and the ‘pacemaker cells’ decrease to uncouple the activity, resulting in reduced frequency during late SNA. Meanwhile, upon the second wave of neurogenesis, more ventrally-located secondary motoneurons and INs are recruited forming a new microcircuit with HF activity containing a higher number of gap junctions with the ‘pacemaker cells’. Fast and slow swimming behavior have been shown to be mediated by a dorsal-ventral topological map of neurons that correspond to the birth order of motoneurons in zebrafish larvae (McLean et al., 2007; McLean & Fetcho, 2009) as well as the molecular identity of motoneurons and V2a interneurons in adult fish (Pallucchi et al., 2024). One intriguing future study would be to test how disturbing the activity of the HF and LF microcircuits during coiling behavior affect fast and slow swimming in zebrafish.

**Figure 8.**
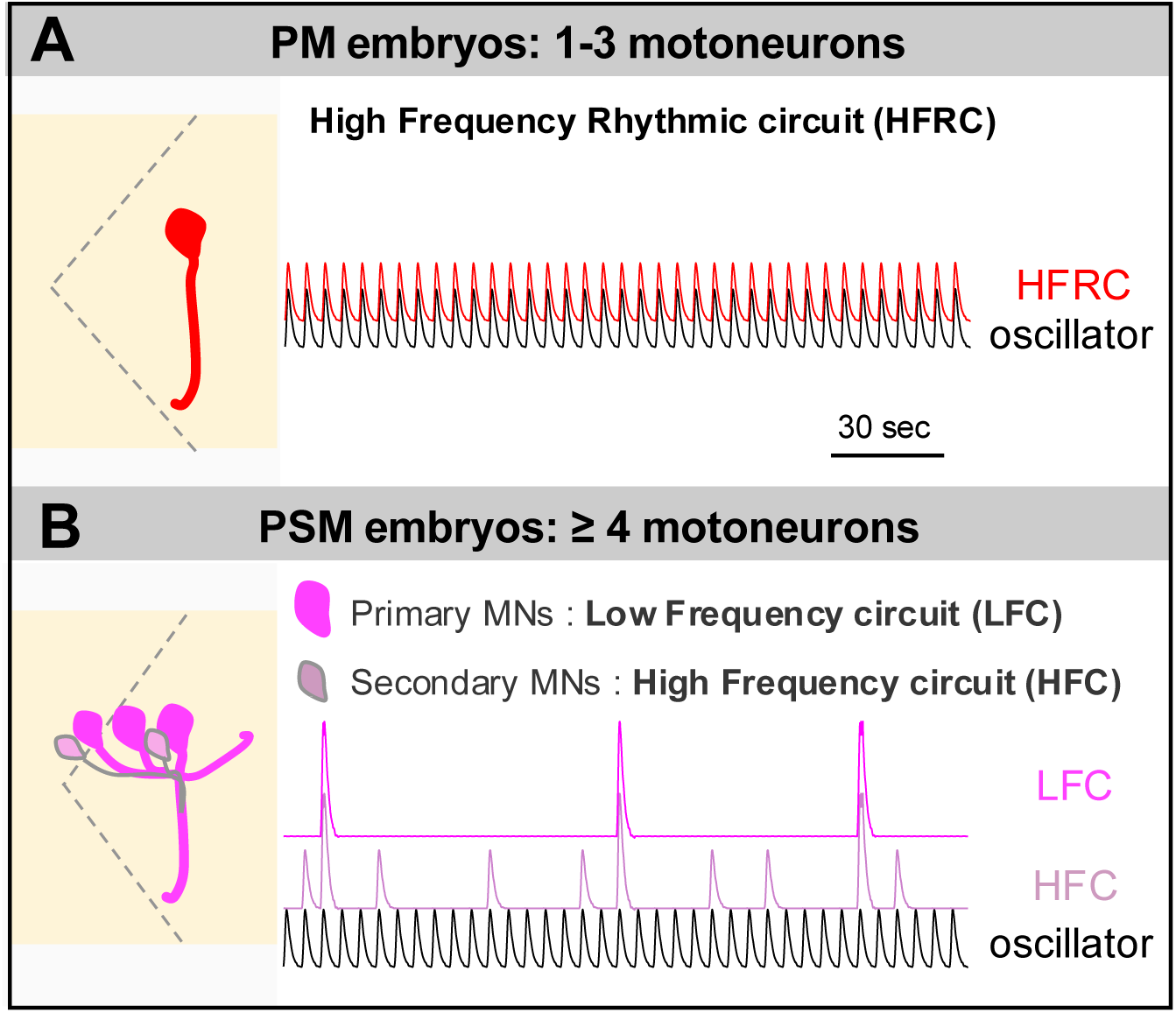
Schematic model for the evolution of spontaneous neural activity (SNA) in zebrafish spinal neurons. (A) Younger embryos expressing mnx1:RFP in only primary motoneurons (1-3 motoneurons (red)) per bundle, display single activity type that is tightly coupled with the oscillator (black). The high frequency rhythmic activity forms a high-frequency rhythmic circuit that control left-right muscle contractions. (B) Older embryos expressing mnx1:RFP in both primary and secondary motoneurons (over 4 motoneurons per bundle), consist of primary motoneurons (magenta) that have uncoupled with the oscillator to display low-frequency (LF) activity forming the low-frequency circuit. Secondary motoneurons (pink) (still lightly coupled with the oscillator) display high-frequency (HF) activity forming the high-frequency circuit. Also see Figure S3.

### Construing left-right muscle contractions by calcium imaging of laterally mounted embryos

Calcium imaging studies mainly analyze signals in the neuron soma. In this study, high resolution imaging and analysis of laterally mounted embryos revealed the presence of puncta and axons that display anti-correlated activity to the ipsilateral neuron soma. Analysis of muscle contractions in unparalyzed embryos show two distinct types of alternating muscle contractions indicative of left-right alternating coiling behaviour in laterally mounted embryos. Combining muscle contraction analysis with calcium imaging of ipsilateral neurons strongly suggest that the signals from the ipsilateral puncta/axons correspond to the contralateral activity. The results show that high resolution calcium imaging in laterally mounted embryos is a powerful method to reconstruct left-right coiling behavior during SNA in the zebrafish embryo.

### Role of cholinergic signaling in maintaining HF rhythmic activity and left-right alternation in early spontaneous neural activity

In contrast to rodent and chick, studies using cholinergic antagonists in the zebrafish embryo failed to inhibit SNA (Saint-Amant & Drapeau, 2001; Warp et al., 2012). This was interpreted to mean that the neurotransmission mechanisms underlying SNA are different in the zebrafish to other vertebrates. Through detailed analysis of neuronal activity in *cholineacetyltransferase-a (chata)* mutant embryos, we found that the mutants exhibit aberrant arrhythmic and asymmetry activity between the ipsi- and contralateral circuit activity in contrast to the highly uniform activity in WT embryos. The results suggest that cholinergic signalling is necessary to maintain symmetry in frequency and left-right alternation during early SNA in PM embryos. Interestingly, mouse *chat* mutants also exhibit decreased frequency in activity as well as defects in left-right and flexor-extensor coordination (Myers et al., 2005). Our results indicate that cholinergic signaling plays a conserved role in modulating symmetric and uniform activity pattern in the embryonic vertebrate spinal cord during left-right alternating SNA.

We propose a model where acetylcholine released from motoneurons activates nAChRs in ascending and/or descending excitatory INs to synchronize the activity between ipsilateral INs and motoneurons. Acetylcholine also activates the inhibitory commissural INs leading to the inhibition of the contralateral motor circuit. The synchronization of activity between these neurons will lead to the strengthening of their synapses to form a functional circuit. So then why would the attenuation of acetylcholine result in aberrant synchronized activity? During SNA, neurons in the spinal cord are extending their axons and dynamically forming and pruning synapses onto their targets (see Figure S3A). Due to the rostral to caudal development of the spinal cord, the types and number of neurons that form synapses onto the motoneuron bundles in each segment are likely different. In other words, one bundle of motoneurons may contain more excitatory synapses, while in the other segment have more inhibitory synapses. Acetylcholine could play a role in allowing yet weakly connected IN-MN synapses to exert their full weight to equalize their influence on the activity of motoneurons during left-right alternation and between between different rostro-caudal segments. For example, in the absence of acetylcholine, the activity would become irregular as neurons that have more inhibitory inputs would have a higher probability to be inhibited during each pacemaker “beat”, resulting in longer intervals between events while neurons with more excitatory inputs would exhibit higher frequency activity. Overall, we propose that cholinergic signaling plays a role in enhancing the uniformity between events and symmetry of left right alternation during early SNA. Future studies should focus on identifying the nAChR subunits as well as muscarinic receptors that are expressed in different IN types.

Where is the source and site of acetylcholine release? To our knowledge, there is no descending cholinergic neurons from the brain at this stage of development in zebrafish (Rima et al., 2020) suggesting that the only neurons that release acetylcholine are the motoneurons. We propose that the rhythmic activity during early SNA results in the release of acetylcholine from motoneurons that is augmented via the activation of α7 nAChRs expressed initially in the primary CaP and later in all motoneurons (Figure S2B). Acetylcholine release could be from two non-mutually exclusive sites: from either the soma of the motoneurons or at the axo-dendritic site between the motoneuron and INs, increasing the strength of IN-MN synapses via presynaptic nAChRs (Rima et al., 2020). In the CNS, acetylcholine is released at non-synaptic sites through volumetric release (Picciotto et al., 2012). Moreover, somatic release of acetylcholine was reported in cultured *Xenopus* motoneurons (Sun & Poo, 1987; Welle et al., 2018; Young, 1986). Future studies using an acetylcholine sensor will help determine the release site for acetylcholine from motoneurons during SNA (Jing et al., 2020).

### Modulation of high frequency activity by cholinergic signaling

nAChR antagonists have a milder effect in the zebrafish compared to rodents (Hong et al., 2013). We instead used nicotine, an agonist of nAChRs, to probe the contribution of cholinergic signaling during SNA. In our model, we propose that the high-frequency rhythmic activity occurring in early PM embryos transitions to LF activity in PSM embryos while the newly born neurons display HF activity. However, the HF activity in PSM embryos is not as uniform in frequency and left-right alternation as during early SNA. The HF activity during late SNA in PSM WT embryos resembles the early activity profile in the *chata* mutants. The result suggests that the HF arrhythmic activity is due to decreased influence and/or attenuation of cholinergic signaling. If so, increasing cholinergic signaling should convert the late HF activity to the early rhythmic HF activity. While we found a general increase in the activity frequency upon application of nicotine, the variability in their response necessitates further studies.

In conclusion, our work illustrates the emergence of birth order determined functional microcircuits and provides support for a conserved role of cholinergic signaling in the vertebrate spinal cord during SNA.

## Acknowledgements

We thank Isaac Bianco and Misha Ahrens for providing transgenic zebrafish lines, Marion Baraban for reagents, Yara Lattouf with fluorescent in situ hybridization and the technical support from the IBPS aquatic and imaging facilities. This study was supported by a graduate fellowship from the Alexis and Anne Marie Habib Foundation (to M.A.Y.) and the Fondation pour la Recherche Médicale (to P.L., J.-M.M. and E.H.).

## Author contributions

M.A.Y., M.R., P.L., J.-M.M., and E.H. designed experiments. M.A.Y., M.R., J.-P.C., E.R., F.H., E.B. and E.H. conducted the experiments. M.A.Y., M.R., N.P., F.H., J.-P.C., and E.H. analyzed data. M.A.Y. and E.H. wrote the manuscript with input from all authors.

## Supplemental Information

**Table S1.**
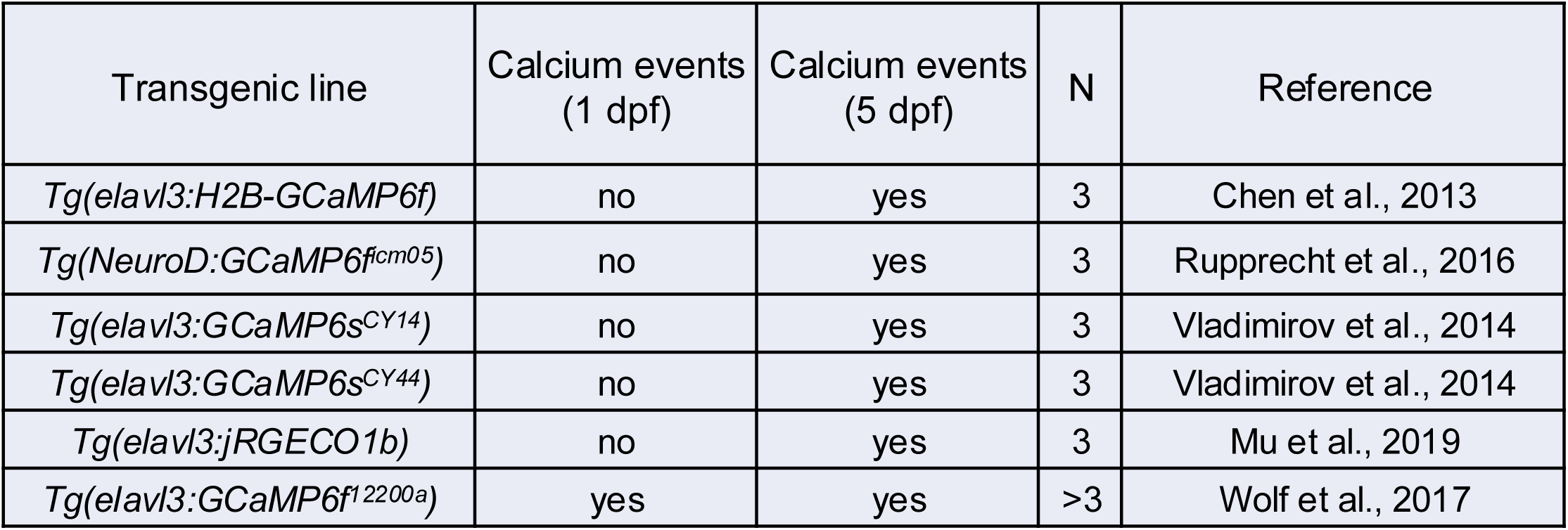
Transgenic lines expressing GCaMP calcium sensors tested for calcium transients in 1-day old embryos.

**Figure S1.**
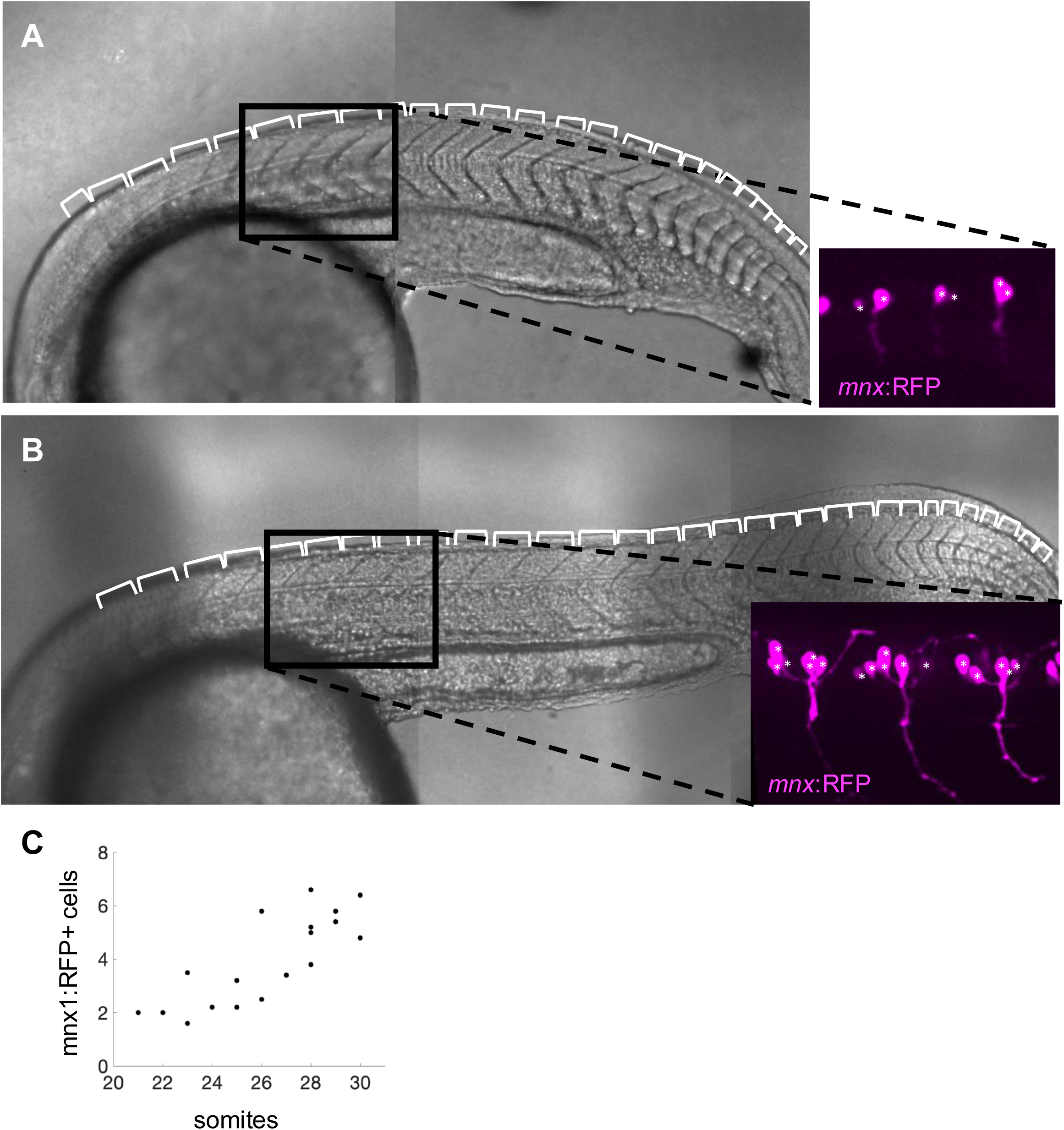
Relationship between somite number and mnx:RFP^+^ cells in the pivot point, related to Figure 3. (A) Transmitted light image of a lateral view of a 25 somite (white brackets) embryo showing 1-2 mnx:RFP^+^ (magenta) cells per bundle. (B) Transmitted light image of a lateral view of a 30 somite (white brackets) embryo showing 5-6 mnx:RFP^+^ (magenta) cells per bundle. (C) Scatter plot showing correlation between somite number and the average number of mnx:RFP+ cells per bundle in the pivot point. N = 19 embryos, n = 89 motoneuron bundles.

**Figure S2.**
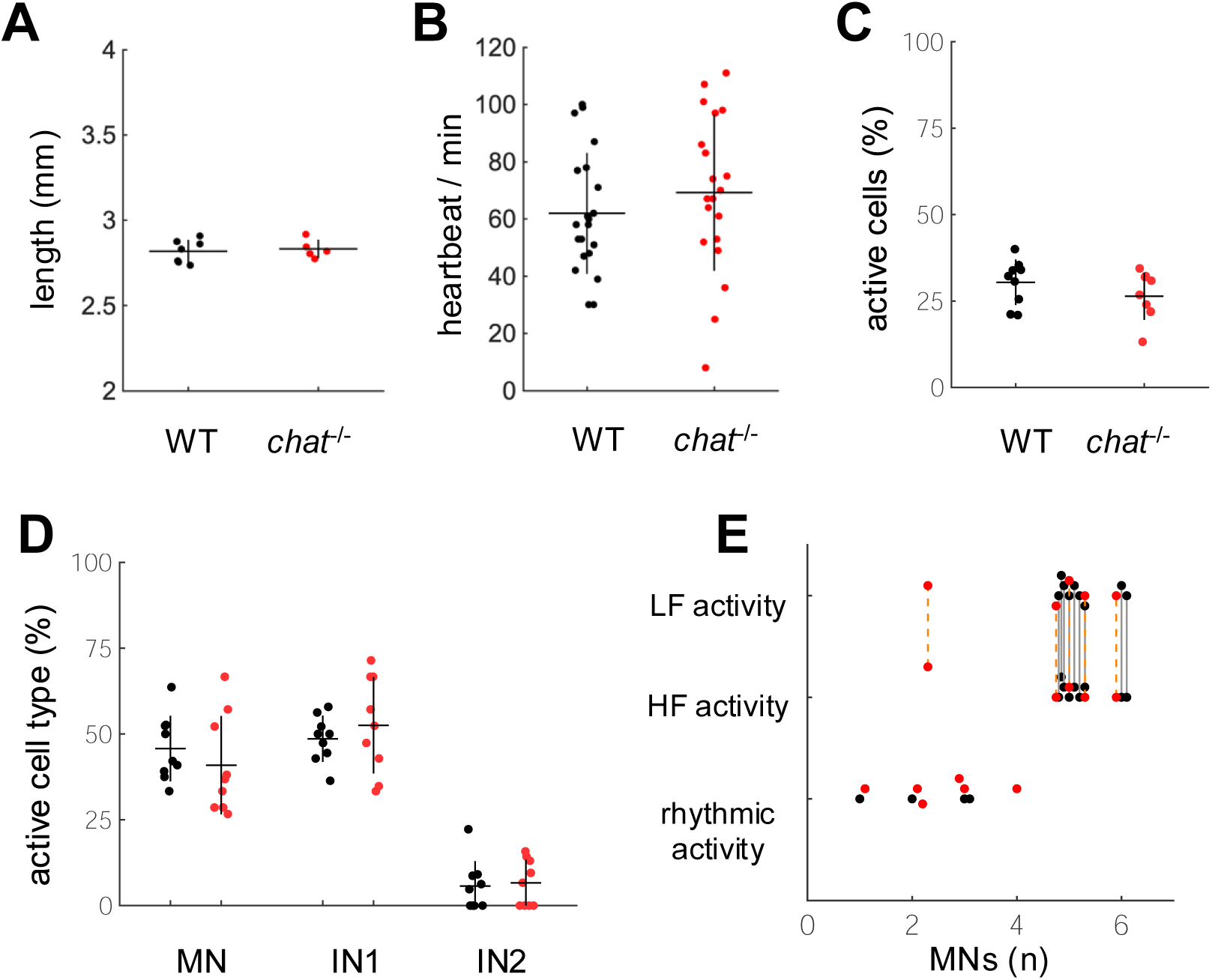
*chata* mutants do not display overt differences with their WT siblings, related to Figure 7. (A) Length of embryos in WT (N = 7) and mutant (N = 5) embryos: 2.82 ± 0.07 vs. 2.83 ± 0.05 µm. (B) Number of heartbeats in 1 min in WT (N = 27) vs. mutant (N = 32) embryos: 54.4 ± 23.9 vs. 55.9 ± 29.1. (C) Scatter plot showing the percentage of active cells in WT and *chata ^-/-^* 1-day old embryos. WT: N = 12, *chata^-/-^ N* = 10, Mann-Whitney test p=0.1915. (D) Scatter plot showing the percentage of neuron types (MN, IN1, & IN2) in active cells between WT and *chata^-/-^* embryos. WT (N = 12) vs. *chata^-/-^* (N= 10); MN: 43 ± 9.5 vs. 44.4 ± 12.21 %, IN1: 46.7 ± 11.1 vs. 46.17 ± 12 %, IN2: 12.9 ± 7.3 vs. 15.6 ± 8 %. (E) Scatter plot of the the distribution of activity types in embryos staged by the number of mnx1:RFP^+^ cells (MN) per bundle. Activity types on the left y-axis correspond to rhythmic activity, high frequency (HF) or low frequency (LF) activity. Older PSM embryos displaying two circuits are represented by a line between HF and LF activity. WT: N = 12, *chata^-/-^* :N = 11.

**Figure S3.**
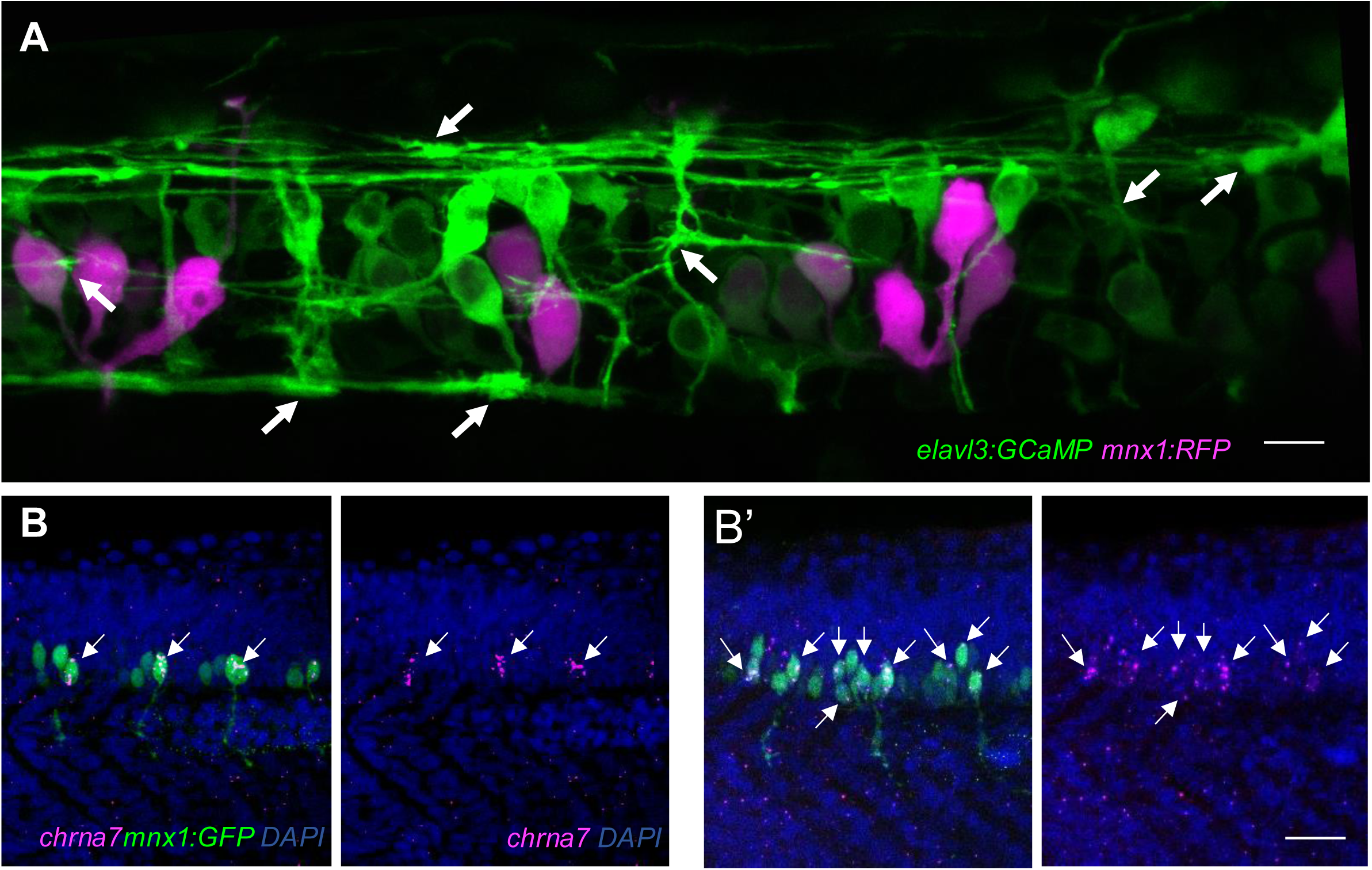
Dynamic changes occur during early embryonic stages, related to Figure 8. (A) Axon elongation and dynamic protrusions occur in many spinal neurons during the evolution of spontaneous neural activity. Z-stack composite confocal image of *Tg(elavl3:GCaMP6f;mnx1:gal4;UAS:RFP*) embryo showing motoneurons (magenta) and spinal neurons (green). White arrows point to putative growth cones. Scale bar = 10 µm. (B,B’) Fluorescent in situ hybridization of *chrna7* (magenta) in *Tg(mnx1:GFP)* (green) embryos. Note that *chrna7* is first expressed only in CaP motoneurons (B) but later becomes expressed in most motoneurons (B’). Arrows indicate *chrna7* labeling. Scale bar = 50 µm.

**Movie S1. Spinal neurons have two functional circuits showing high frequency (red) vs. low frequency (yellow) activity.**

**Movie S2. WT PM embryos show regular alternation in calcium activity between motoneuron soma (green asterisks) and in puncta (purple arrow).**

**Movie S3. *chata* mutant PM embryos show irregular alternation in calcium activity between motoneuron soma (green asterisks) and in puncta (purple arrow).**

---

**Video S1. Spinal neurons have two functional circuits showing high frequency (red) vs. low frequency (yellow) activity, related to Figure 3.**

**Video S2. WT PM embryos show regular alternation in calcium activity between motoneuron soma (green asterisks) and in puncta (purple arrow), related to Figure 7.**

**Video S3. *chat* mutant PM embryos show irregular alternation in calcium activity between motoneuron soma (green asterisks) and in puncta (purple arrow), related to Figure 7.**

